# When and why spatial structure favors metabolic cross-feeding

**DOI:** 10.64898/2026.10.09.757969

**Authors:** Jian Wang, Ihab Hashem, Satyajeet Bhonsale, Jan F.M. Van Impe

## Abstract

Metabolic cross-feeding is central to microbial life. Theory predicts that spatial structure favors cross-feeding by affording cells preferential access to the exchanged metabolites. Yet recent evidence shows that spatial structure can, in some contexts, disfavor cross-feeding. Here, we resolve this contradiction by structuring cross-feeding interactions into three dimensions: (i) directionality, (ii) investment, and (iii) metabolic dependency. We then investigated their combined effects using a spatially explicit individual-based model. We find that the effect of spatial structure on cross-feeding is determined not by physical proximity alone, but by how microbial growth depends on exchanged metabolites. Metabolic dependence shapes the spatial organization of cells and metabolite fluxes, and this emergent organization, in turn, reshapes the ecological outcome of cross-feeding. Our results reconcile previously conflicting observations and provide a general framework for predicting and engineering metabolically interdependent microbial communities.

**Significance Statement:** Cross-feeding, the sharing of metabolites among microbes, is one of the most common interactions in nature, and it helps build the communities that drive processes ranging from gut digestion and crop production to global nutrient cycling. A long-standing view says that physical association among microbes should promote this sharing. Here, we show this view is incomplete. Whether spatial structure strengthens or weakens cross-feeding depends on the type of metabolic dependence between the partners, not on the physical association alone. This finding reconciles conflicting experimental and theoretical observations reported over the past decade and provides a predictive framework for understanding metabolite exchange in natural microbial communities.

**Graphical Abstract:** 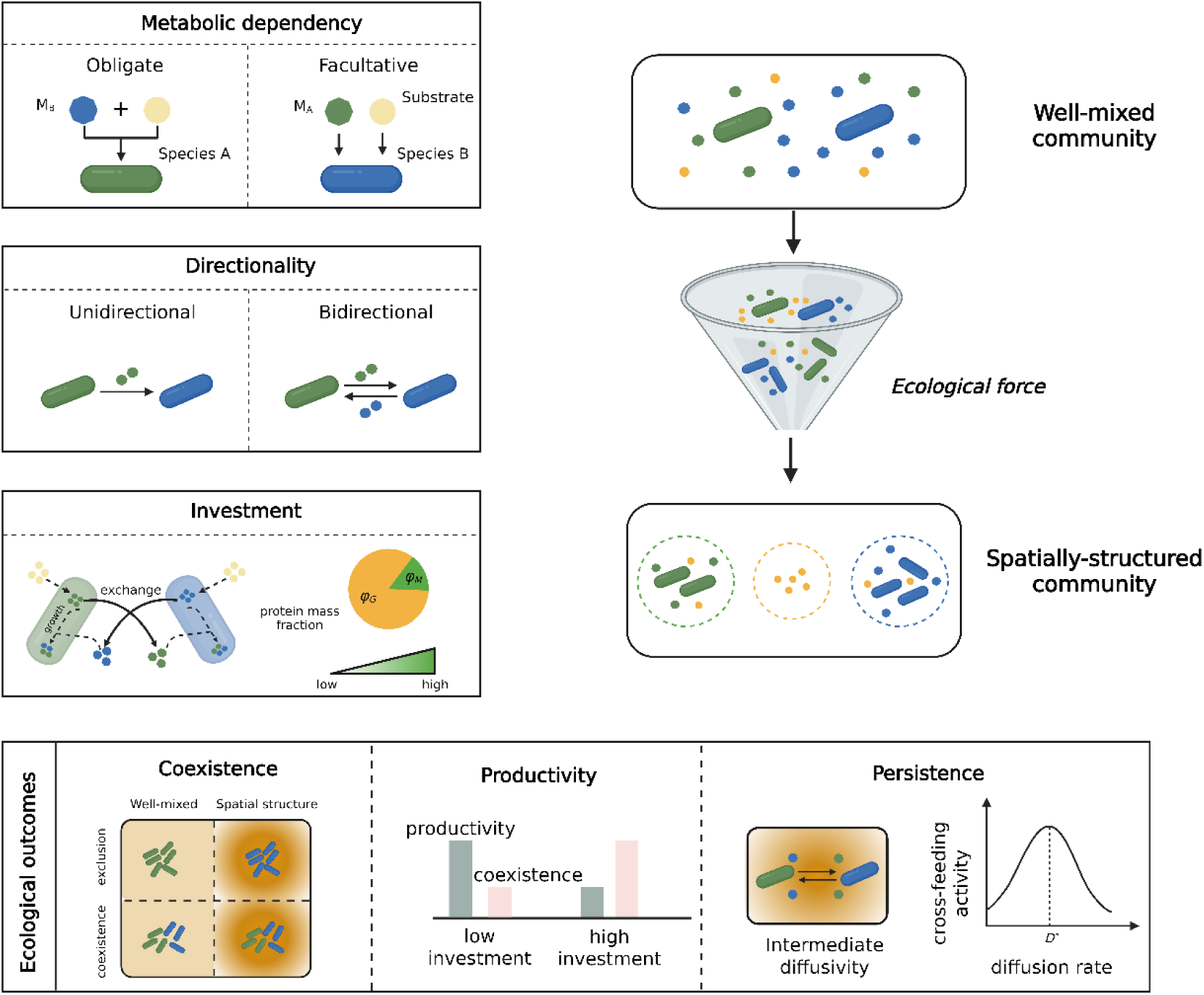

## Introduction

Microbial species rarely make everything they need (1–3). Comparative genome surveys show that many bacteria lack one or more pathways for synthesizing essential metabolites, such as amino acids, vitamins, and even nucleotides (4–7). Consequently, these species depend on metabolites supplied by other community members to sustain growth, a phenomenon known as metabolic cross-feeding (8–10). The evolutionary process likely to drive the emergence of cross-feeding is called the Black Queen Hypothesis (BQH) (11). The BQH predicts that selection can favor the loss of costly, leaky functions when neighboring species reliably supply their products. Such adaptive gene loss creates robust ecological links among community members (12–14). Over the past two decades, sequencing and co-occurrence analyses have revealed widespread metabolic cross-feeding in microbial communities (15–17). Studies have shown that metabolic cross-feeding shapes community composition (18–20), stabilizes coexistence (21–23), and expands collective metabolic niches (24–26). Understanding the ecological and evolutionary conditions that favor cross-feeding is therefore essential for predicting and controlling microbial communities.

In natural populations, cross-feeding is inherently spatial because metabolites must diffuse to recipient cells before being diluted, degraded, or transported away (27, 28). Spatial structure localizes both microbial cells and the metabolites they release, making secreted products more likely to be taken up by nearby partners than dispersed throughout the wider community. This localization generates positive assortment between producers and the recipients of their metabolites. When recipients provide reciprocal or indirect benefits, such assortment can direct returns toward producers while limiting access by nonproducing *‘hitchhikers’* (29). Spatial structure has therefore long been viewed as an important mechanism for maintaining costly metabolite exchange and promoting its evolution (30–33).

However, a growing body of literature shows that localization does not necessarily favor cross-feeding. Using a multispecies biofilm model, Mitri et al. (2011) showed that conditions increasing assortment within species can simultaneously segregate mutualistic partners, thus preventing costly secretors from efficiently receiving return benefits (34). Chen et al. (2023) found that the assembly of a metabolically specialized consortium under diffusion limitation was governed more by metabolic flux than by spatial proximity (35). Lobanov et al. (2023) further showed that slow metabolite diffusion can weaken interaction strengths and reduce microbial coexistence (36). Especially, Luo et al. (2024) argued that spatial structure can suppress cooperative behavior by prolonging cooperators’ exposure to nearby *‘hitchhikers’*. Therefore, whether spatial structure favors or disfavors cooperative metabolic interactions remains debated. On the one hand, limited dispersal can increase local metabolite concentrations and direct exchanged metabolites toward nearby partners. On the other hand, the same spatial constraints can separate complementary partners, restrict metabolite transport, and intensify local competition for shared resources. The balance between these opposing effects depends on how cross-feeding is metabolically organized (37) .

D’Souza et al. (2018) proposed a conceptual framework to help understand the origin and evolution of metabolic cross-feeding (8). They classify cross-feeding along two dimensions: (i) the directionality of metabolite flow (i.e., unidirectional versus bidirectional), and (ii) the investment made by producers in supplying exchanged metabolites. This two-dimensional framework describes who supplies metabolites to whom and at what cost. However, they do not specify the degree to which receiver growth depends on the exchanged metabolite. We therefore introduce metabolic dependency as a third dimension (i.e., obligate versus facultative). Obligate receivers require the exchanged metabolite for growth, whereas facultative receivers obtain only an additional growth benefit from it. We argue this dimension is important because metabolic dependency determines how local metabolite availability translates into microbial growth and, consequently, how interaction dynamics both shape and respond to spatial organization.

Here, we study metabolic cross-feeding interactions in a minimal two-species microbial consortium. We incorporate a wide range of cross-feeding interactions (i.e., varying metabolite-exchange direction, producer investment, and metabolic dependency) into a spatially explicit individual-based model (IbM). We compare their ecological outcomes in well-mixed and spatially structured environments. We find that metabolic dependency, the way receiver species use metabolites, shapes community spatial organization and determines whether spatial structure promotes or suppresses cross-feeding. Under open boundaries, we further show that cross-feeding efficacy is maximized at an intermediate metabolite diffusivity, which balances transport to receivers with metabolite degradation and environmental loss. Together, these conceptual and modeling results provide a mechanistic basis for predicting when, and explaining why, spatial structure shapes the outcome of metabolic cross-feeding.

## Results

### A three-dimensional framework for disentangling metabolic cross-feeding

Cross-feeding interactions can be classified along three axes: (i) directionality, whether metabolic exchange flows from one species to another (i.e., unidirectional) or is reciprocated (i.e., bidirectional); (ii) investment, the fraction of a producer’s metabolic flux diverted from its own growth to metabolite secretion; and (iii) dependency, whether a receiver requires the cross-fed metabolite to grow (i.e., obligate) or only benefits from it (i.e., facultative). Depending on how the receiver uses the metabolite, facultative cross-feeding can be further classified as sequential, parallel, or synergistic. These dimensions give a wide range of cross-feeding interactions (Fig. 1). We implemented all of them in an IbM framework to investigate how spatial structure shapes community properties.

**Fig. 1.**
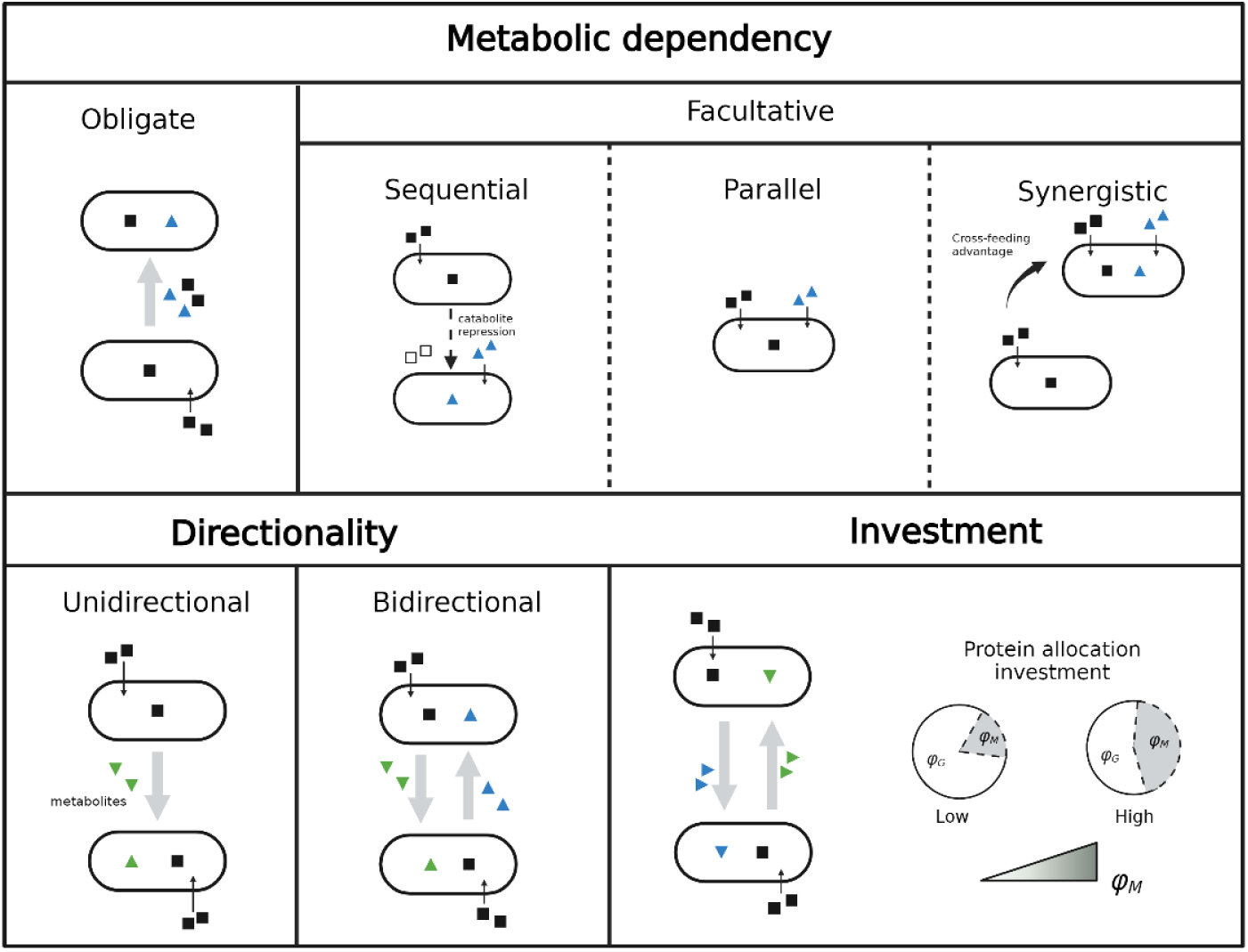
Types of metabolic cross-feeding interactions. Cross-feeding interactions are classified along three axes: (i) metabolic dependency, whether the receiver requires the exchanged metabolite for growth (i.e., obligate) or only benefits from it (i.e., facultative), (ii) directionality, whether metabolic flows from one species to another (i.e., unidirectional) or is reciprocated (i.e., bidirectional), and (iii) investment, the fraction of a producer’s protein mass allocation diverted from its own growth to metabolite secretion (quantified by *φ_M_*). This classification extends the framework of D’Souza et al. (2018) by incorporating metabolic dependency and treating investment as a continuous variable (8).

IbM represents each microbial cell as a discrete entity with explicit growth kinetics, metabolite secretion, and physical interactions (38, 39) (Fig. 2A-C). Cells grow on a shared substrate according to Monod kinetics and divert a fraction of *I* of their gross metabolic capacity to metabolite secretion (40). The net specific growth rate of species is

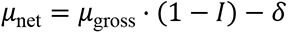

**Fig. 2.**
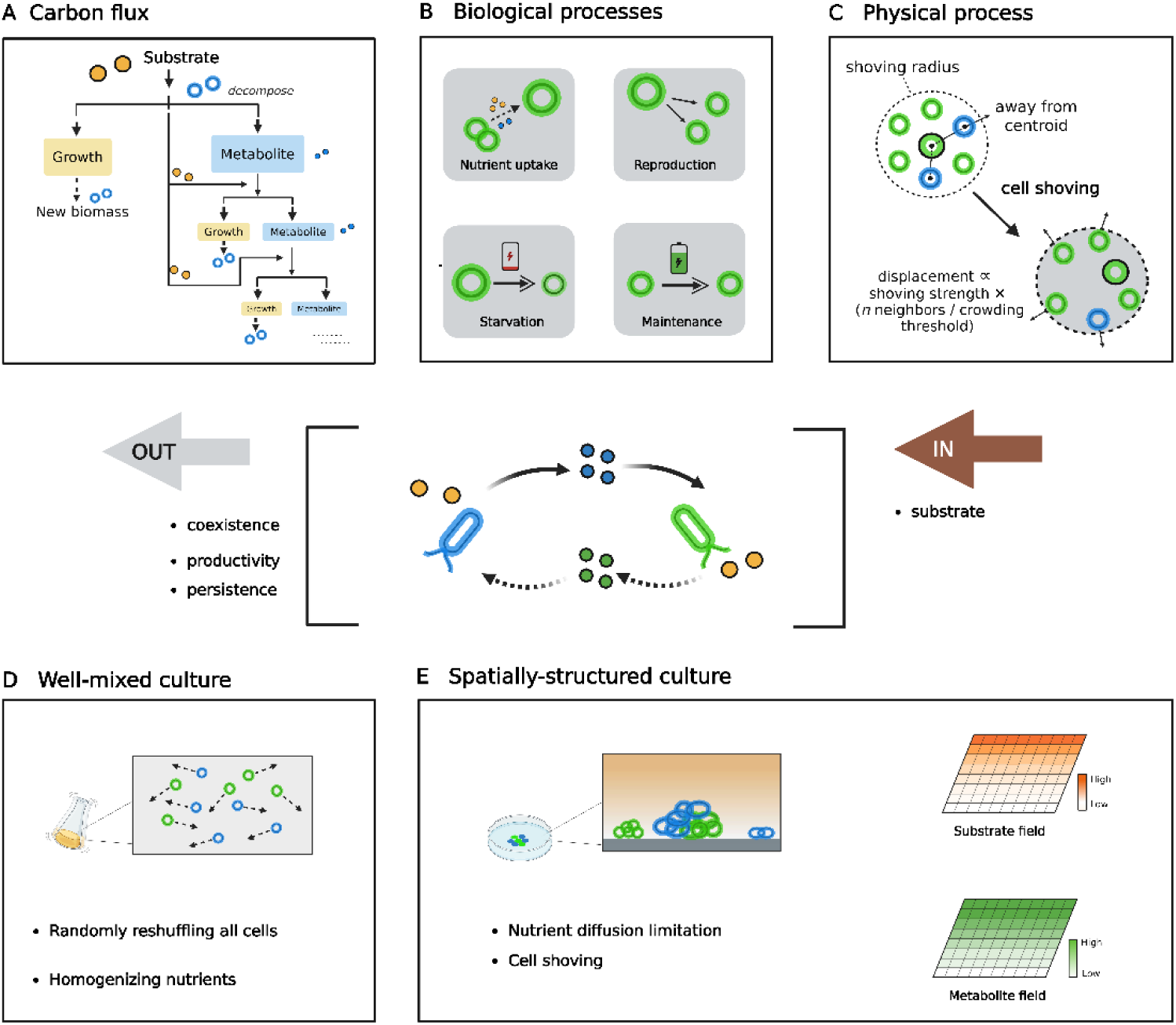
Overview of the individual-based model (IbM) for a range of metabolic cross-feeding interactions. (A) Carbon flux pathways. Two species convert a shared primary substrate into biomass and metabolites. The exchanged metabolites support partner growth through cross-feeding. (B) Cell-level biological processes. Individual cells take up available nutrients to meet maintenance requirements, grow, divide, and secrete metabolites. Cells enter a starvation state when nutrient uptake is insufficient to satisfy maintenance demands. (C) Physical interactions. A cell-shoving algorithm resolves mechanical interactions and prevents spatial overlap between growing cells. Simulations were conducted under two environmental conditions. (D) In well-mixed cultures, cell positions are randomly reshuffled, and nutrient concentrations are spatially homogenized at each time step to represent shaken liquid cultures. (E) In spatially structured cultures, cells remain fixed on the surface, while the primary substrate and exchanged metabolites diffuse according to Fick’s law.

With *μ*_gross_ [1/h] the gross growth rate on substrate and metabolite (Methods, *Growth rate equations*) and *δ* [1/h] the maintenance rate. Here, we focus on a minimal consortium where two species engage in cross-feeding mediated by diffusible metabolites. Species *A* secrets metabolite *M_A_* and species *B* secrets metabolite *M_B_*. Both metabolites diffuse with coefficient *D*. In all scenarios below, both species A and B invest in secretion at the same level, regardless of directionality. In bidirectional cross-feeding, species *A* uptake metabolite *M_B_*, and species *B* uptake metabolite *M_A_*. In unidirectional cross-feeding, species *B* still invest in secreting its metabolite, even though this metabolite is not taken up by species *A*. This model specification is to remove the effect of unequal investment between species and to ensure that the unidirectional receiver remains metabolically contributing rather than acting as a nonproducing ‘*hitchhiker’*. We simulated consortium growth in two cultures: (i) a well-mixed culture, in which cell positions are randomly reshuffled, and nutrient concentrations (i.e., substrate and metabolites) are homogenized at each time period (Fig. 2D), and (ii) a spatially structured culture, in which diffusion is limited so that individuals are in local interactions with their neighbors (Fig. 2E). Each simulation was initialized with the two species inoculated at equal abundance in a random configuration (Methods, *Inoculation*). Communities were expanded until the primary substrate was depleted to below 1% of initial concentration. We used coexistence and community productivity (i.e., total abundance of species *A* and *B*) to indicate the outcome of these interactions (Methods, *Statistics*).

### Investment stabilizes coexistence at the cost of community productivity in obligate consortia

We first examined obligate cross-feeding, in which the receiver requires the cross-fed metabolite to grow. For obligate bidirectional cross-feeding, also known as mutualism, both partners had to exceed a minimum investment threshold before the interaction could be initiated. Above this threshold, coexistence remains high due to strict mutual dependency. This *‘all-or-nothing’* coexistence pattern is generated by an ecological Allee effect (41) (Fig. 3A-B). This pattern is also consistent with previous observations that engineered microbial mutualisms converge toward an intermediate species ratio across a wide range of initial frequencies (42). Under well-mixed conditions, however, the coexistence index declined measurably (i.e., from nearly 1 to 0.9) as both species increased their investment in parallel. Under spatial structure, coexistence stayed high across the same investment range (Fig. 3C). Examination of species abundances shows that the species with a high intrinsic growth rate gradually expands its advantage over its partner. Spatial structure confines this competitive advantage to local neighborhoods and prevents the faster-growing species from dominating the community (Fig. S3).

**Fig. 3.**
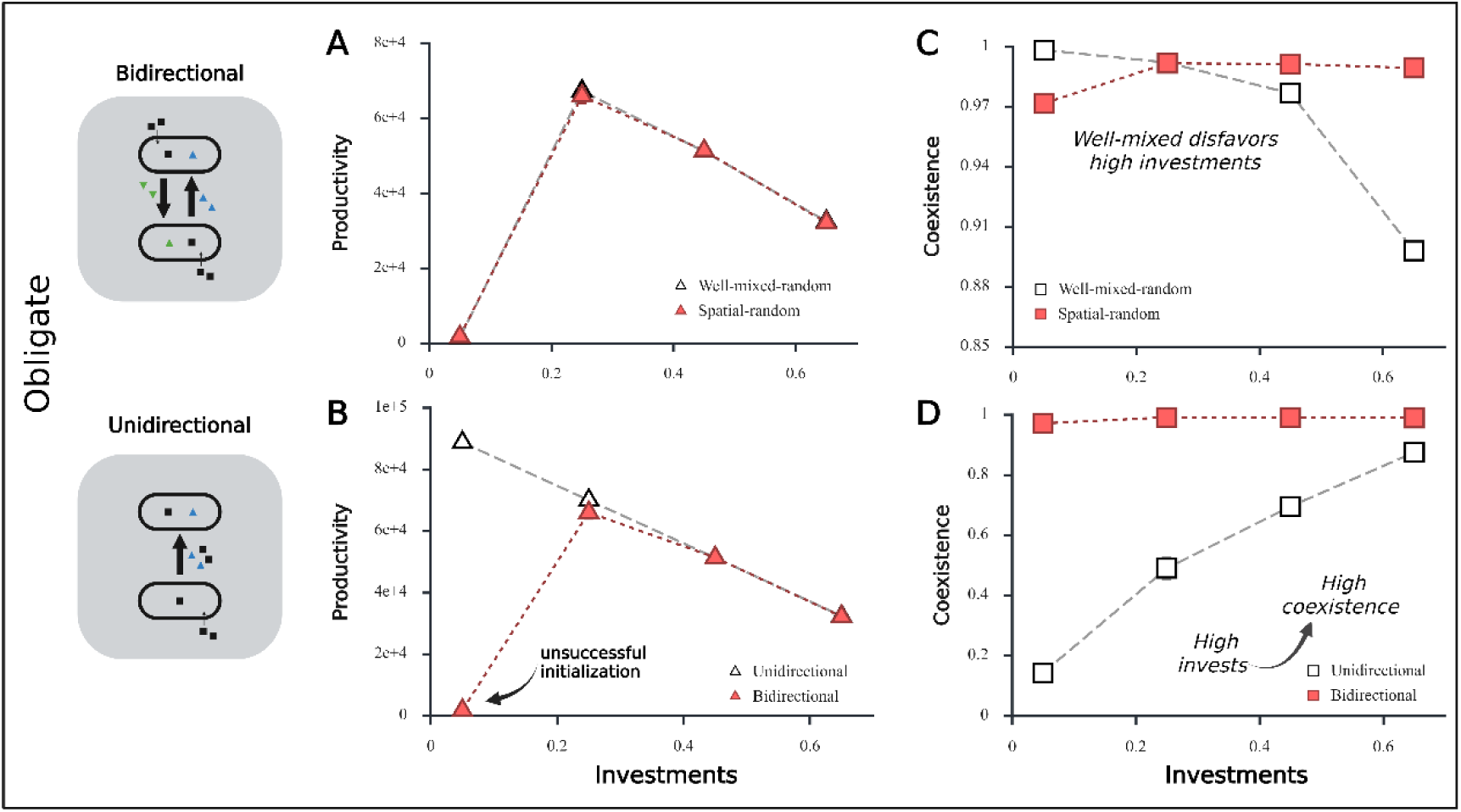
Investment governs coexistence in obligate cross-feeding consortia. (A) Initiation threshold for obligate bidirectional cross-feeding. Under both well-mixed and spatially structured conditions, bidirectional cross-feeding is initiated only when both species invest above a minimum level. Below this level, metabolite supply is insufficient to sustain partner growth, and the consortium collapses. (B) Obligate unidirectional cross-feeding can be initiated even at low producer investment because the producer grows independently on the primary substrate, and cumulative metabolite secretion eventually supports receiver growth. (C) In bidirectional consortia, coexistence remains high across the investment range (∼0.9). Under well-mixed conditions, however, increasing investment slightly reduces coexistence by amplifying the growth disadvantage of the slower-growing species (from ∼1 to ∼0.9). Spatial structure limits the expansion of the faster-growing species and maintains a consistently high coexistence index (∼ 0.97). (D) In unidirectional consortia, increasing producer investment strengthens metabolite supply to the receiver while reducing the producer’s own growth capacity. These opposing effects reduce abundance asymmetry and promote coexistence, although spatial structure prevents excessive receiver expansion at the producer’s expense.

We then examined obligate unidirectional cross-feeding, in which species *A* secretes a metabolite that species *B* strictly require without reciprocation. This one-way benefit flow lets the receiver exploit the producer at no cost to itself, also known as parasitism. Unlike the bidirectional system, unidirectional cross-feeding could be initiated even at low producer investment (Fig. 3B). Coexistence nevertheless remained low because metabolite supply was insufficient to sustain receiver growth. As investment increases, metabolite availability rises and, with the decrease in the producer’s growth potential, the abundance of the two species becomes more even. Under well-mixed conditions, higher investment sharply reduced producer abundance and eventually reversed species’ dominance (i.e., the receiver came to dominate the community). Meanwhile, as the producer declined, total metabolite supply also fell, and receiver abundance subsequently decreased. Spatial structure prevented this reversal. Local retention of the secreted metabolite limited receiver expansion and protected the producer from competitive exclusion. Spatial structure therefore allowed the consortium to sustain higher levels of costly investment without losing the producer (Fig. 3D). This finding is consistent with the general principle that spatial localization can protect costly producers from exploitation by noncontributing partners (34, 43).

### Metabolic dependency determines whether spatial structure promotes or suppresses coexistence

We next asked whether metabolic dependency changes the consortium’s response to spatial structure. We compared obligate unidirectional cross-feeding with facultative unidirectional cross-feeding. In the facultative scenario, the receiver can grow on substrate alone, and the cross-fed metabolite provides an additional benefit. Under well-mixed conditions, facultative receivers gained two competitive advantages. They have higher intrinsic growth rates and have uniform access to both the substrate and producer-derived metabolites. Consequently, receivers outcompeted producers, and the coexistence index remained low (i.e., around 0.2). Spatial structure weakens receivers’ advantage by localizing the producer-derived metabolites. Only receivers near producers obtained the full metabolic benefit, which limited their expansion across the community and reduced the risk of producer exclusion. Spatial structure therefore increased coexistence in facultative consortia. Fig. 4A shows that the coexistence index increased by approximately 93.8% under spatial structure conditions compared with well-mixed conditions.

**Fig. 4.**
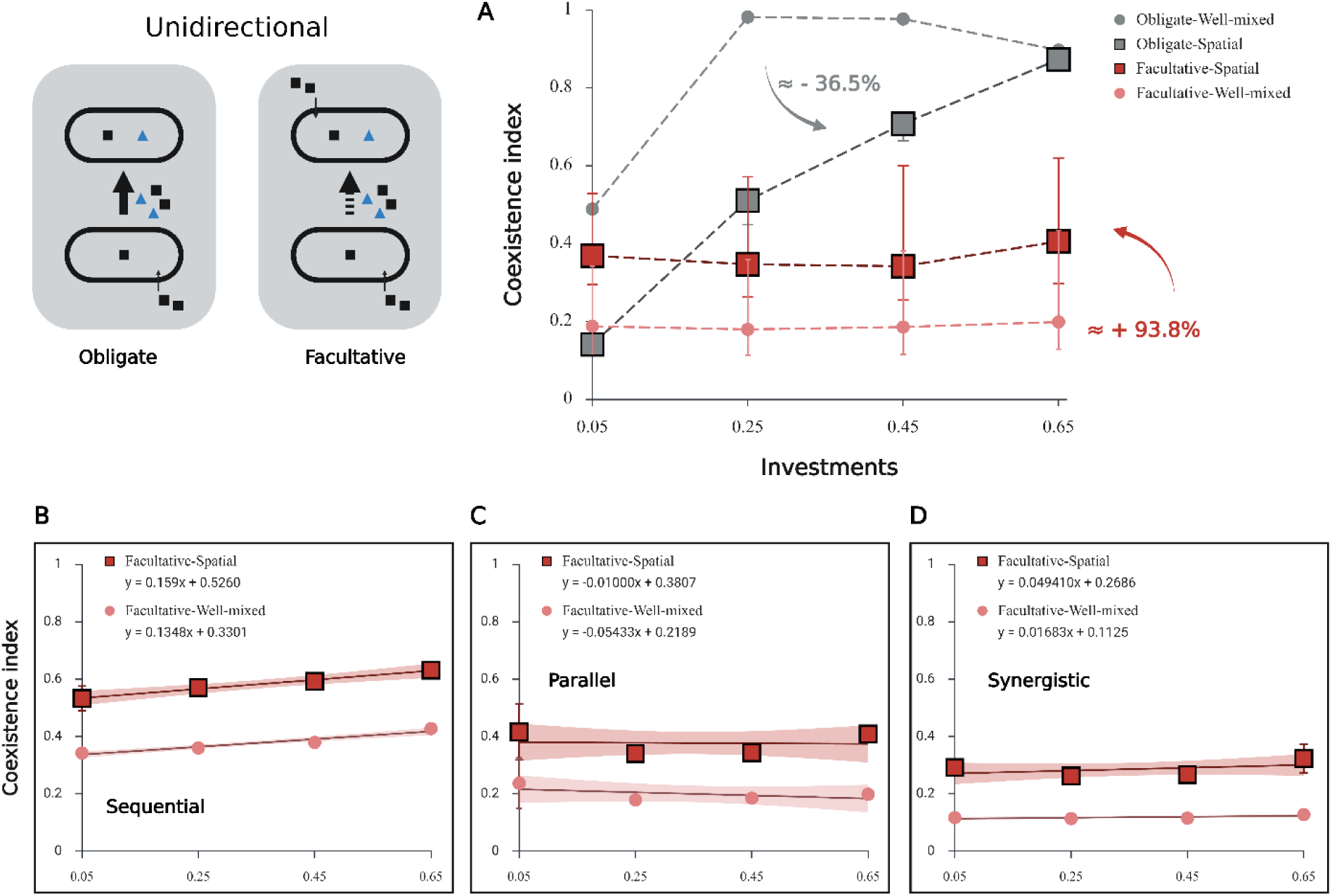
Metabolic dependency determines the sign of spatial effects on coexistence. (A) Coexistence index for obligate and facultative unidirectional cross-feeding, under well-mixed and spatially structured conditions. Spatial structure increases coexistence in facultative consortia by 93.8% relative to the corresponding well-mixed condition because metabolite localization restricts the additional competitive advantage of receivers to cells located near producers. By contrast, spatial structure decreases coexistence in obligate consortia by 36.5% because spatially segregated receivers lose access to a metabolite required for growth, which weakens the metabolic coupling between partners. Symbols represent mean values from three independent simulation replicates (n = 3). Error bars indicate ±1 standard deviation (SD). (B–D) Coexistence index as a function of investment for the three facultative sub-types, sequential (B), parallel (C), and synergistic (D), under well-mixed and spatially structured conditions, with linear fits; facultative consortia maintain a comparatively stable coexistence index across the investment range under both conditions. Shaded bands in B–D indicate 95% confidence intervals for the fitted mean.

The opposite pattern appears in obligate consortia. Obligate receivers cannot grow without the producer-derived metabolite. Well-mixed conditions distribute metabolites throughout the environment and allow receivers to uptake them regardless of position. By contrast, spatial structure confined metabolites to the producer neighborhood and reduced their availability to spatially segregated receivers. Spatial structure therefore weakens the metabolic coupling required for receiver persistence and reduces coexistence by approximately 36.5% relative to well-mixed conditions. Interestingly, the same physical process of local metabolite retention gives rise to opposite ecological outcomes under different forms of metabolic dependency. These results challenge the traditional view that spatial structure favors cross-feeding and show instead that metabolic dependency determines whether spatial structure stabilizes or destabilizes coexistence of cross-feeding consortia.

Metabolic dependency also altered how consortia responded to investment. In obligate consortia, increasing investment had two opposing effects on the receiver. Higher producer investment increased metabolite supply and reduced producer growth, which weakened competition and initially allowed the receiver population to expand. However, at higher investment levels, the receiver was subject to the same investment cost as the producer (both species were specified to invest equally in metabolite secretion even for unidirectional cross-feeding). The resulting loss of growth capacity eventually outweighed the benefit of increased metabolite supply, and receiver abundance declined. Facultative consortia responded differently because receivers could grow on the primary substrate without the exchanged metabolite. Increased metabolite supply therefore provided only an additional benefit and did not release the receiver from an absolute growth limitation. As investment increased, both species allocated more metabolic capacity to secretion, and their biomasses declined at similar rates. Their relative abundances consequently changed little, even though total community biomass decreased. Accordingly, facultative consortia maintained a comparatively stable coexistence index across the investment range under both well-mixed and spatially structured conditions (Fig. 4B–D).

### Spatial structure favors slower-growing species at high investment

Having established that metabolic dependency regulates the effects of spatial structure on coexistence, we next asked whether spatial structure benefits both partners equally in bidirectional cross-feeding. We compared all bidirectional interaction types under well-mixed and spatially structured conditions.

In bidirectional cross-feeding, each species acts as both a producer and a receiver. Differences in intrinsic growth rate naturally create an initial advantage for the faster-growing species. The magnitude of this advantage depends on metabolic dependency. Fig. 5A shows that the slower-growing species gains increased advantages under a higher investment level. The slower-growing species had the lowest advantage in obligate consortia because strict mutual dependence constrained the two partners to grow together (Fig. 5B). It had the highest advantage in facultative-sequential consortia, where growth depended mainly on competition for the shared substrate (Fig. 5C).

**Fig. 5.**
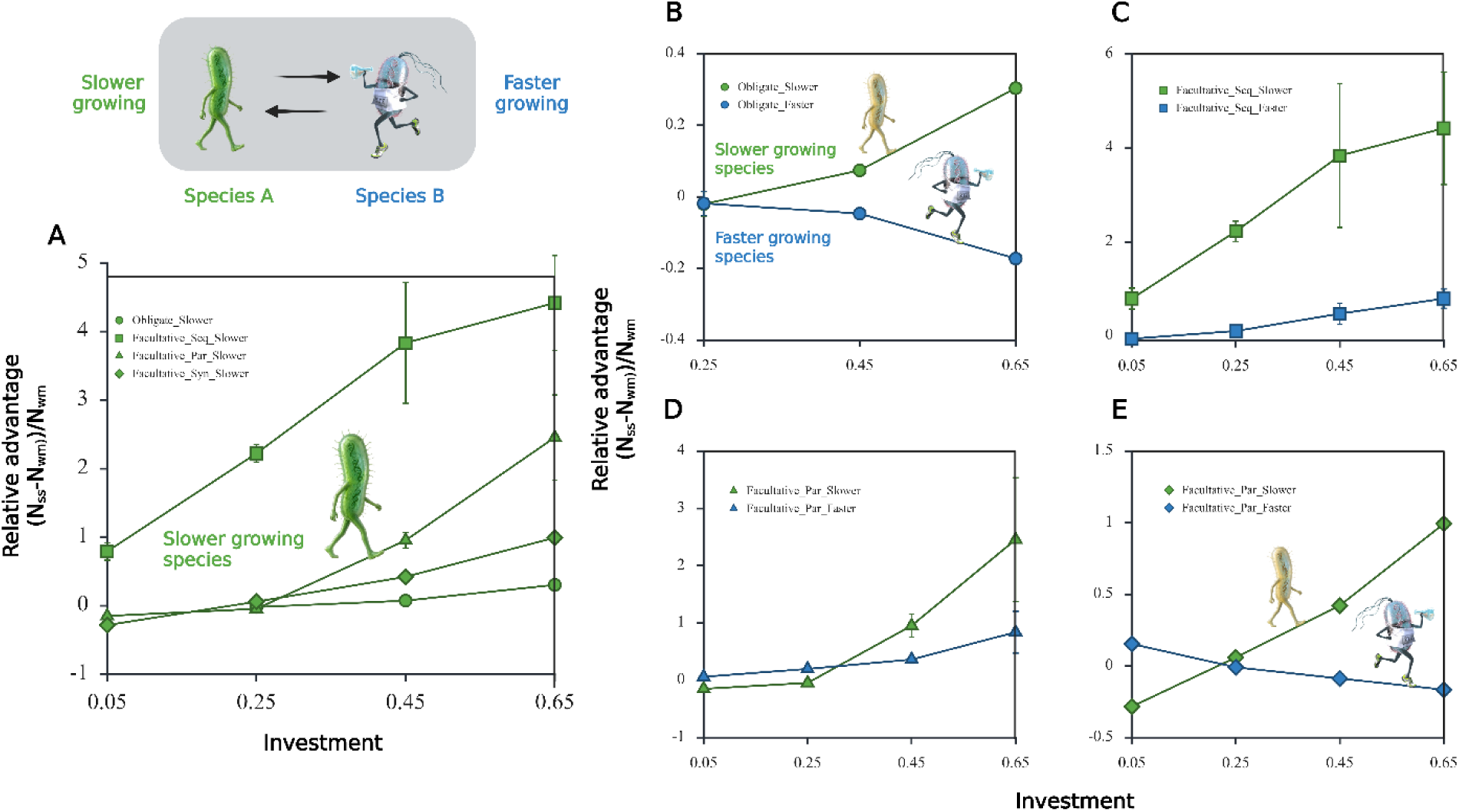
Spatial structure favors the slower-growing species at high investment in bidirectional cross-feeding. (A) Across all dependency types, spatial structure amplifies the relative advantage of slower-growing species in the consortia. Relative advantage calculated by (*N*_spatial_ − *N*_well-mixed_)/*N*_well-mixed_. (B-E) Relative effects of spatial structure on the slower- and faster-growing species for all dependency types: obligate (B), facultative sequential (C), facultative parallel (D), facultative synergistic (E). At low investment, competition for the shared substrate dominates, and localized substrate depletion strengthens the faster-growing species’ advantage. At high investment, partner-derived metabolites contribute more strongly to growth, and their local retention compensates for the slower-growing species’ competitive disadvantage. Spatial structure therefore increasingly favors the slower-growing partner as investment rises. Symbols represent mean values from three independent simulation replicates (n = 3). Error bars indicate ±1 standard deviation (SD). The cartoons depicting the slower- and faster-growing species were adapted from Li et al. (2019), with artwork credited to Victor Leshyk (44).

We found that the effect of spatial structure on the slower-growing species changed with investment. At low investment, both species secreted little metabolite, so cross-feeding provided only a weak benefit. Direct competition for the shared substrate remained the dominant interaction. Under spatial structure, the faster-growing species depleted substrate within its local neighborhood and reduced substrate availability to nearby cells of the slower-growing species. The limited metabolite supply was insufficient to compensate for this competitive local disadvantage. Well-mixed conditions distributed the substrate uniformly and prevented such localized depletion, which allowed the slower-growing species to maintain a higher abundance.

At high investment, both species allocated a larger fraction of their metabolic flux to secretion, and metabolite exchange became more important relative to direct substrate competition. The performance of each species increasingly depended on access to metabolites supplied by its partner. Spatial structure retained these metabolites within producer neighborhoods and created locally enriched exchange zones. The slower-growing species therefore received a reliable metabolic benefit from nearby faster-growing partners. Under well-mixed conditions, metabolite concentrations were homogenized across the population, so the slower-growing species did not experience the same local enrichment, while the faster-growing species retained its intrinsic growth advantage. Spatial structure therefore benefited the slower-growing species most strongly at high investment because local metabolite access increasingly compensated for its disadvantage in substrate competition. Ecologically, it shows that spatial structure can favor slower-growing species at high investment and help maintain a more balanced community.

### An intermediate diffusivity maximizes cross-feeding under open conditions

We next asked whether cross-feeding can persist when exchanged metabolites are lost from the local environment. The simulations above used closed (periodic) boundaries, which conserve all metabolites within the domain. Natural microbial habitats, however, are rarely closed. Metabolites can diffuse out of soil pores, biofilms, and host-associated microhabitats, or disappear through chemical and biological degradation. Cross-feeding therefore depends not only on local metabolite retention, but also on whether metabolites can reach their partners before they are lost.

We examined this transport constraint in obligate bidirectional consortia. We replaced the periodic boundaries with absorbing boundaries, where any metabolite that reaches the edge of the domain is removed from the system. We then varied metabolite diffusivity *D* and decay rate *δ* to determine the conditions under which cross-feeding could be initiated and maintained. We hypothesized that diffusivity would have a non-monotonic effect on promoting cross-feeding. At low diffusivity, metabolites remain concentrated near producers but may decay before reaching spatially segregated partners. At high diffusivity, metabolites diffuse farther but are rapidly dispersed and lost through the open boundaries. Their concentration at receiver locations can then fall below the threshold required for growth. These opposing constraints predict that there should exist an intermediate diffusivity at which metabolites diffuse far enough to reach their partners while remaining sufficiently concentrated to support growth. To visualize the metabolic exchange effects clearly, we tested this hypothesis using a separated inoculation geometry. Each species occupied one half of a domain of length *L*, with absorbing boundaries at the outer edges. The simulations confirmed an intermediate optimum (Fig. 6D). Cross-feeding failed at low *D* because metabolite transport between partners was insufficient. It also failed at high *D* because rapid dispersal and environmental loss reduced metabolite concentrations below the activation threshold. Coexistence persisted only across an intermediate range of diffusivity.

**Fig. 6.**
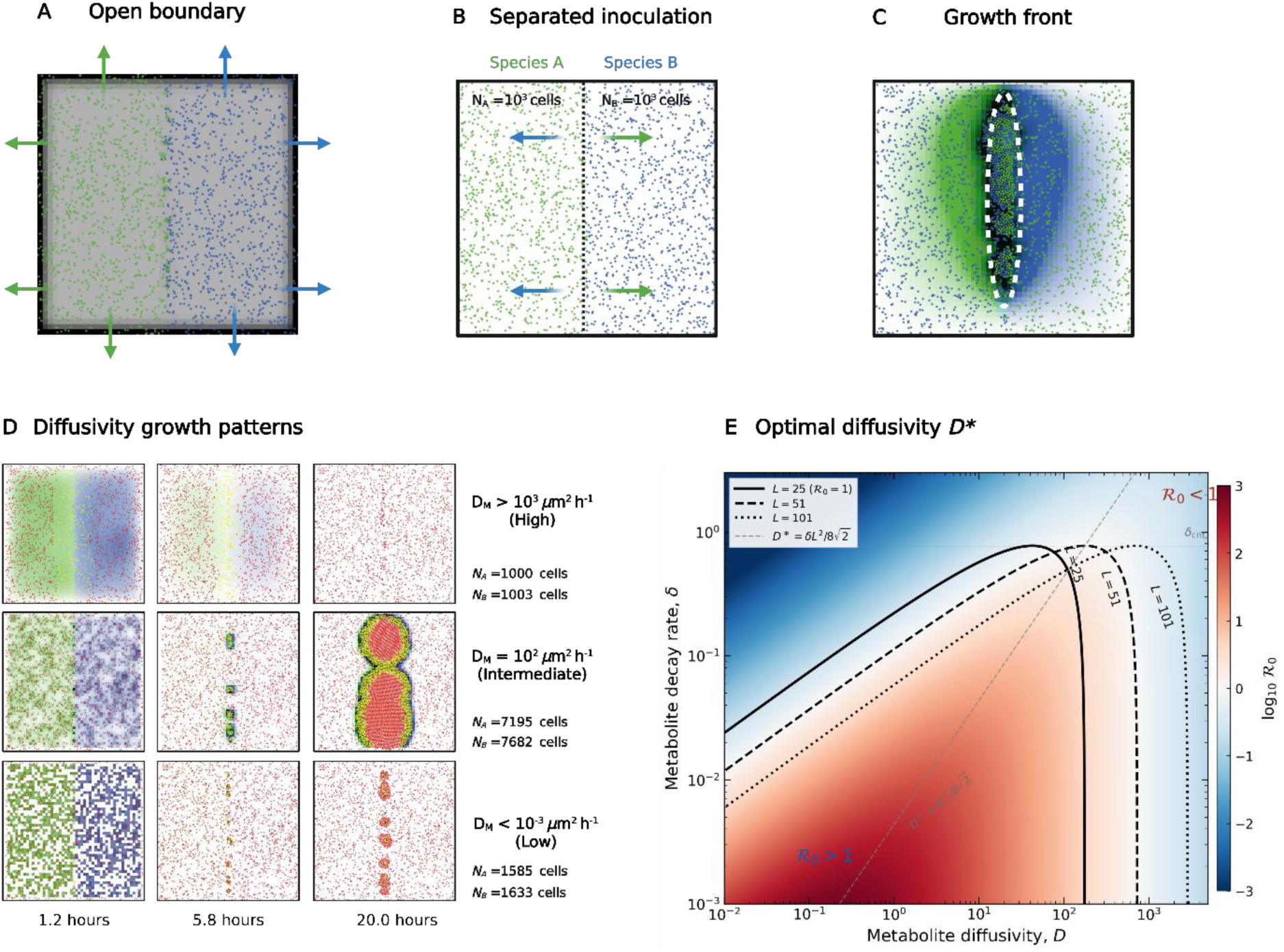
| An intermediate metabolite diffusivity maximizes cross-feeding under open boundaries. (A) Open (absorbing) boundary condition, in which a metabolite reaching the domain edge is removed from the system. (B) Separated inoculation geometry, with each species occupying one half of a domain of length L. (C) Growth front of an obligate bidirectional cross-feeding consortium, with reciprocal metabolite exchange supporting growth at the interface between the two species (white dashed outline). (D) Representative spatial snapshots of growth at low, intermediate, and high metabolite diffusivity, with final population sizes of species A and B and the corresponding simulated time; cross-feeding is most productive at intermediate diffusivity. (E) Analytical derivation of cross-feeding reproduction number R_0_ as a function of metabolite diffusivity D, showing a single interior maximum that defines the optimal diffusivity D*.

We summarized this tradeoff analytically by solving the steady-state reaction–diffusion equation for the metabolite field and coupling metabolite availability to receiver growth through Monod kinetics (SI, methods). This analysis yielded a dimensionless cross-feeding reproduction number, *R*_0_. Cross-feeding persisted when *R*_0_ > 1and collapsed when *R*_0_ < 1. This threshold resembles the investment threshold observed in obligate bidirectional consortia, but here it is determined by metabolite transport rather than metabolic allocation. The reproduction number contains two opposing components. A transport term increases with diffusivity because metabolites are more likely to reach distant partners. A local-retention term decreases with diffusivity because metabolites are more rapidly dispersed and lost. Their product reaches a single maximum and therefore defines an optimal diffusivity, *D*^∗^. Ecologically, this result shows that neither maximal retention nor maximal dispersal is optimal for cross-feeding. Metabolite transport must instead match the distance between partners and the rate of environmental loss.

## Discussion

Natural microbial communities are often thought of as a cruel place, with most species ruthlessly competing for space or resources as they struggle to survive (45, 46). However, a meta-analysis of 77 studies showed that metabolite cross-feeding is widespread among naturally occurring microbes (8, 47). More evidence indicates that metabolic cross-feeding is essential for structuring the composition, assembly, and functionalities of microbial communities across diverse habitats (10, 48–50). Understanding the ecological and evolutionary conditions that allow such positive interactions to emerge and persist has therefore become a central question in microbial ecology (51, 52).

Much of our mechanistic understanding of cross-feeding has been developed in trackable experimental systems (53). Engineered consortia, defined growth media, and microfluidic devices have made it possible to manipulate cell spacing, metabolite supply, and interaction strength with high precision. These approaches have revealed key mechanisms underlying the emergence of metabolic exchange and have provided practical insights into the control of cross-feeding communities (26, 32, 51, 54) . However, natural environments expose microbial interactions to a wider range of biotic and abiotic constraints (55). Nutrients are often scarce, and interacting species continue to compete for common resources even as they supply metabolites to one another.

Spatial structure is particularly important because it reshapes the relationship between metabolite exchange and resource competition (56–58). Under well-mixed conditions, exchanged metabolites and shared resources are distributed across the entire community. The benefits of cross-feeding and the costs of resource competition are experienced at the same population scale. In contrast, spatial structure confines both processes to local neighborhoods. Nearby cells become the principal competitors for shared substrates, but they are also the most likely recipients of exchanged metabolites. Therefore, the central question has become how local metabolite exchange interplays with local resource competition and how this interplay is translated into microbial growth.

Here, we classify metabolic cross-feeding along three axes: (i) directionality, (ii) investment, and (iii) metabolic dependency and incorporate these interactions into a spatially explicit individual-based model. By simulating microbial consortia growth in both well-mixed and spatially structured environments, our simulations reveal four main findings. First, investment defines the boundary between the establishment and collapse of obligate cross-feeding. Higher investment increases metabolite supply, but it also diverts metabolic capacity away from producer growth. Spatial structure can afford producer species with higher investment by limiting receiver expansion at the expense of producers. Second, metabolic dependency determines the sign of spatial effects on cross-feeding. Spatial structure favors facultative cross-feeding because localization restricts the additional competitive benefit obtained by receivers. The same localization disfavors obligate cross-feeding because it deprives spatially segregated receivers of a metabolite required for growth. Third, investment overturns the dominance between community members. Competition for the shared substrate governs community organization at low investment, whereas metabolic coupling becomes increasingly important at high investment. Spatial structure can consequently shift from favoring intrinsically faster competitors to supporting slower-growing partners that benefit from locally retained metabolites. Finally, metabolite transport imposes a physical limit on these spatial effects. Under open boundaries, cross-feeding is maximized at an intermediate metabolite diffusivity by balancing transport to receivers with metabolite degradation and environmental loss Our framework makes several simplifying assumptions that warrant further investigation. We consider only ecological interactions among partners and do not model the evolutionary dynamics of investment or secretion efficiency. In natural communities, investment is subject to selection. Depending on the ecological context, selection may favor either reduced investment and the emergence of nonproducing hitchhikers or greater investment and stronger cooperation. Extending the model to include heritable variation in investment and producer–hitchhiker dynamics would allow us to determine whether the coexistence patterns identified here remain stable over evolutionary timescales. We also represent exchanged metabolites without specifying their biochemical identities. Incorporating metabolite-specific diffusivity, uptake kinetics, stoichiometric constraints, and degradation rates would improve quantitative predictions for particular cross-feeding systems (59–61). Additionally, our coexistence index measures the balance between species abundances at the end of a single growth cycle. Such finite-time coexistence can be transient and does not establish long-term stability (62). Whether both species persist across repeated growth cycles remains to be tested.

Overall, our results highlight the importance of metabolite dependency in regulating the spatial effects of cross-feeding. Our conceptual and modeling framework provides a tractable and generalizable approach to inform the design of synthetic microbial consortia in engineered environments.

## Methods

### Growth rate equations

We performed simulations in the individual-based modeling toolkit MICRODIMS.logo (*In Review*), which is a derived version of MICRODIMS (63, 64). A central design feature is the explicit separation of gross and net specific growth. The gross rate captures intrinsic substrate- and metabolite-driven kinetics, *μ*_gross_ = *μ*_max_ · *S*/(*K_S_* + *S*) · *g*(*M*), with *g*(*M*) = 1 for cells acting solely as producers under obligate dependency.

The net rate *μ*_net_ = *μ*_gross_ · (1 − *I*) − *δ* imposes an explicit growth–secretion tradeoff through the investment fraction *I* ∈ [0,1] and a maintenance drag δ. Secretion flux = *φ* · *λ* · *μ*_gross_ · *m* · *Δt*, with secretion efficiency *λ* scaling production independently of the net growth cost. Diverse metabolic interactions follow the unified growth equation (65–67):

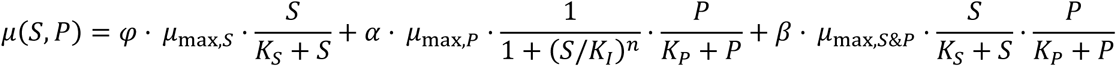

with each dependency type corresponding to a specific weight set (SI Appendix, Table 1). Cell mass accumulates as *m*(*t* + *Δt*) = *m*(*t*)(1 + *μ*_net_*Δt*) with *μ*_net_ ≥ 0; on reaching a division threshold, the cell halves and hatches a daughter at a small random offset (forced positive in biofilm mode). Intraspecific heterogeneity is introduced by drawing each cell’s *μ*_max_ at birth within ±*ε* of the species mean. Physical crowding is resolved by an iterative pressure-based shoving algorithm.

### Diffusion

Substrate and metabolite transport uses the NetLogo diffuse operator; the physical diffusivity is mapped to a dimensionless fraction *d* = min(0.5, max(0, *DΔt*/*Δx*²)), applied in three substeps per tick, followed by first-order metabolite decay *c_M_*(*t* + *Δt*) = *c_M_*(*t*)(1 − *kdΔt*). Three boundary regimes implement distinct environments: well-mixed (all patches replaced by their global mean each tick, cells randomly repositioned), colony (free diffusion with emergent local gradients; optional chemostat replenishment), and biofilm (Dirichlet substrate boundary at the top row, no-flux below, upward-biased division). Closed simulations use periodic boundaries; open simulations use absorbing boundaries. Details and the seven inoculation modes appear in SI Appendix, S4.

### Community-level analytics

Coexistence is quantified by *φ* = 4 · *N_A_* · *N_B_*/(*N_A_* + *N_B_*)² ∈ [0,1] (SI Appendix, S1). Productivity is the total final biomass *N* = *N_A_* + *N_B_*.

## Data and Code Availability

All data and code used to replicate the figures, analyses, and findings in this paper can be found at: https://gitlab.kuleuven.be/u0154531/metabolic-cross-feeding.git

## Author Contributions

All authors conceptualized the study. J.W., I.H., and S.B. conducted the investigation. J.W. and I.H. designed the model. J.W. performed the simulation and wrote the manuscript. I.H. and J.F.M.V.I. supervised the work. J.F.M.V.I. provided funding support. All authors edited the manuscript and approved the final version.

## Acknowledgments

The authors thank Francesco Moro, Prof. Bas Teusink, and Prof. Matti Gralka (VU Amsterdam) for their useful discussions.

## Additional Information

Supplementary information can be found in the supplementary material.

