## Supplementary material for "When and why spatial structure favors metabolic cross-feeding"

### Electronic supplementary material

This supplementary material provides the mathematical and computational formulation of the spatial cross-feeding model. The following sections present S1, the model scope and simulation design; S2, growth, metabolic dependency and metabolite exchange; S3, diffusion and environmental loss; S4, outcome metrics; S5, model parameters; S6, the metabolite-ignition threshold; and S7, the reaction–diffusion derivation.

### S1. Model setup and simulation design

The individual-based model represents two microbial species, $A$ and $B$, on a $51 \times 51$ patch lattice. Each cell carries its species identity, mass, maximum specific growth rate and metabolite-production investment. Each patch stores the local concentration of the shared primary substrate $S$ and the two potentially exchanged metabolites $M_A$ and $M_B$. Species $A$ produces $M_A$ and species $B$ produces $M_B$.

The simulated interaction is defined by three properties. Metabolite-flow direction is bidirectional if $A$ consumes $M_B$ and $B$ consumes $M_A$, and unidirectional if only $B$ consumes $M_A$. Metabolic dependency specifies the receiver growth response as obligate, facultative sequential, facultative parallel or facultative synergistic (section S2). Metabolite-production investment $I \in [0, 1]$ specifies the fraction of gross growth capacity diverted to secretion. Both species carry the same imposed value of $I$, including in the unidirectional treatment.

For the comparisons in main-text figures 3–5, both species were inoculated at equal abundance in random positions. Closed simulations used periodic boundaries. In the well-mixed treatment, chemical concentrations were replaced by their domain-wide means and cells were randomly repositioned at every tick. In the spatial treatment, cell positions were retained and chemical fields were redistributed locally by diffusion. The transport analysis in main-text figure 6 used a separated inoculation and absorbing metabolite boundaries (section S3). Simulations ended after the mean primary-substrate concentration fell below 1% of its initial value.

Within each tick, the model (i) evaluates cell growth, substrate uptake, metabolite uptake and secretion from the local chemical fields; (ii) updates diffusion, metabolite decay and the selected boundary condition; (iii) updates cell mass and division; and (iv) resolves local overlap by iterative shoving. The tick duration is $\Delta t = 0.05$ h.

### S2. Growth, metabolic dependency and metabolite exchange

Define the substrate and metabolite response functions

$$f_S(S) = \frac{S}{K_S + S}, \qquad H(M) = \frac{M^n}{\theta^n + M^n}, \qquad f_M(M) = \frac{M}{K_M + M} H(M), \tag{1}$$

and the sequential-use switch

$$q_{\text{seq}}(S) = \frac{1}{1 + \exp[4(S - S^{\ddagger})]}. \tag{2}$$

For a cell wired to consume metabolite $M$, the four implemented gross-growth laws can be written exactly as

$$\mu_{\text{gross}} = \mu_{\max} \left[ w_S f_S(S) + w_M q(S) f_M(M) + w_{SM} f_S(S) f_M(M) \right], \tag{3}$$

with the coefficients in table S1. A cell that is not wired to consume a metabolite uses $\mu_{\text{gross}} = \mu_{\max} f_S(S)$, irrespective of the dependency assigned to its partner.

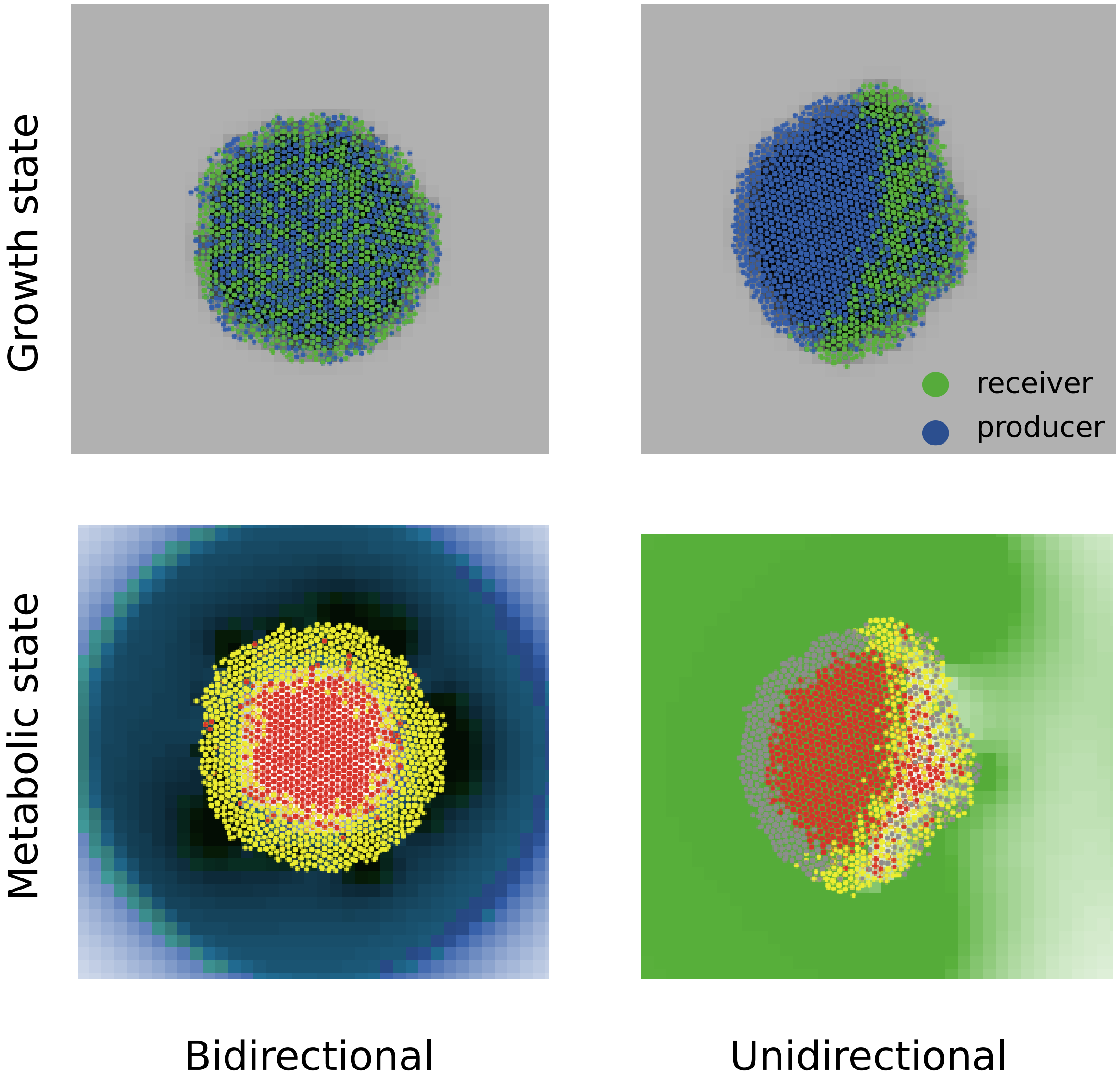


Figure S1: **Cross-feeding directionality controls species intermixing and the spatial layout of metabolic states.** Colony snapshots when substrate is below 90% initial concentration. Left, bidirectional exchange. Right, unidirectional exchange from producer to receiver. Top row, growth state. Cells coloured by species, blue for producer, green for receiver. The grey background is the primary substrate, with darker shades indicating lower concentration. Bottom row, metabolic state. Cells recolored by metabolic mode, grey for growth on primary substrate, yellow for active cross-feeding, red for starved. Patch color shows the secreted metabolites, each drawn in the color of the species that consumes it. Green is the metabolite taken up by the receiver, blue the metabolite taken up by the producer. Saturation increases with concentration, so white indicates depletion.

Table S1: Exact coefficient mapping for the implemented metabolic dependencies in equation 3. A dash denotes a term that is inactive.

| Dependency | $w_S$ | $w_M$ | $q(S)$ | $w_{SM}$ |
|---|---|---|---|---|
| Obligate | 0 | 0 | – | 1 |
| Facultative sequential | 1 | 1 | $q_{\mathrm{seq}}(S)$ | 0 |
| Facultative parallel | 0.5 | 0.5 | 1 | 0 |
| Facultative synergistic | 1 | 0 | – | $\beta_{\mathrm{syn}}$ |

Equation 3 gives, respectively,

$$\mu_{\mathrm{gross}}^{\mathrm{obl}} = \mu_{\max} f_S f_M, \tag{4}$$

$$\mu_{\mathrm{gross}}^{\mathrm{seq}} = \mu_{\max} \left( f_S + q_{\mathrm{seq}} f_M \right), \tag{5}$$

$$\mu_{\mathrm{gross}}^{\mathrm{par}} = \mu_{\max} \left( 0.5 f_S + 0.5 f_M \right), \tag{6}$$

$$\mu_{\mathrm{gross}}^{\mathrm{syn}} = \mu_{\max} f_S \left( 1 + \beta_{\mathrm{syn}} f_M \right). \tag{7}$$

The parallel formulation gives an isolated receiver with $M = 0$ one-half of its maximum substrate-supported rate. This property should be interpreted as part of the implemented parameterization rather than as a generic property of facultative metabolism.

Metabolite-production investment is applied after gross growth is calculated,

$$\mu_{\mathrm{net}} = \max[0, \mu_{\mathrm{gross}}(1 - I) - \delta], \tag{8}$$

where $\delta$ is the maintenance rate. Cell mass is updated as

$$m(t + \Delta t) = m(t) \left[ 1 + \mu_{\mathrm{net}} \Delta t \right]. \tag{9}$$

A cell divides on reaching the division mass and transfers one-half of its mass to a daughter placed at a small random offset. Maximum growth rate is drawn at birth within $\pm 20\%$ of the species mean. Local crowding is resolved through iterative displacement away from the centroid of neighbouring cells.

The metabolite secreted by a producer during one tick is

$$\Delta M_{\text{sec}} = I\lambda\mu_{\text{gross}} m\Delta t, \tag{10}$$

where $\lambda$ scales secretion independently of its imposed growth cost. Substrate and metabolite uptake are

$$\Delta S_{\text{up}} = \min\left(\frac{\mu_{\text{gross}} m\Delta t}{Y_{X/S}},\, 0.8S\right), \tag{11}$$

$$\Delta M_{\text{up}} = \min\left(\frac{\mu_{\max}}{Y_{X/M}}\frac{M}{K_M + M} m\Delta t,\, 0.8M\right). \tag{12}$$

Metabolite uptake in equation 12 is not multiplied by the Hill gate $H(M)$. This difference between production-dependent growth and uptake is central to the ignition threshold derived in section S6.

### S3. Nutrient diffusion

The physical diffusivity $D$ is mapped to the NetLogo diffusion fraction

$$d = \min\left(0.5, \max\left[0, \frac{D\Delta t}{\Delta x^2}\right]\right), \tag{13}$$

and the diffusion update is applied in three substeps per tick. Metabolite decay follows

$$c_M(t + \Delta t) = c_M(t)(1 - k_d\Delta t). \tag{14}$$

Closed simulations retain the periodic boundaries of the wrapped lattice. In the open treatment, metabolite concentrations on the domain edge are reset to zero after diffusion, so metabolites reaching an edge are removed. The well-mixed treatment homogenizes chemical fields and randomizes cell positions at every tick; the spatial treatment retains local concentration gradients and cell positions.

### S4. Coexistence metrics

Let $N_A$ and $N_B$ be the final numbers of living cells. Their relative frequencies and endpoint community evenness are

$$p_A = \frac{N_A}{N_A + N_B}, \qquad p_B = \frac{N_B}{N_A + N_B}, \qquad \phi = \frac{1 - (p_A^2 + p_B^2)}{1 - 1/2} = 4 p_A p_B. \tag{15}$$

The index ranges from zero after loss of one species to one at equal final abundance. It measures endpoint abundance balance, not mutual invasibility or stable long-term coexistence. Joint persistence requires $N_A > 0$ and $N_B > 0$ at the substrate-depletion endpoint. Total final cell abundance is

$$N_{\text{tot}} = N_A + N_B. \tag{16}$$

If summed cell mass is analyzed instead, it should be reported separately as $B_{\text{tot}} = \sum_i m_i$ rather than being conflated with $N_{\text{tot}}$.

The population-specific response to spatial structure in main-text figure 5 is

$$R_i = \frac{N_{i,\text{spatial}} - N_{i,\text{mixed}}}{N_{i,\text{mixed}}}, \qquad i \in \{A, B\}. \tag{17}$$

The percentage effect of spatial structure on endpoint evenness is calculated consistently as

$$100 \times \frac{\phi_{\text{spatial}} - \phi_{\text{mixed}}}{\phi_{\text{mixed}}}. \tag{18}$$

### S5. Parameters used in the reported simulations

Table S2 contains the defaults stated in the supplied model description. Metabolite-production investment, metabolite diffusivity and metabolite decay were varied in the analyses identified in the main text.

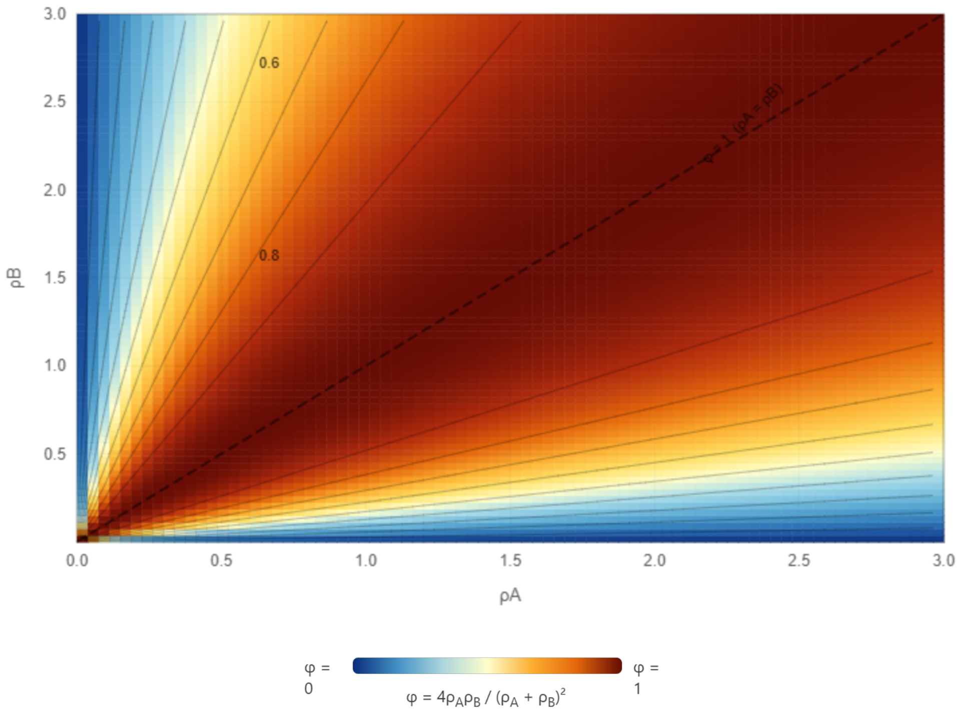


Figure S2: **Geometry of the coexistence index $\varphi$.** Contour map of $\varphi = 4\rho_A\rho_B/(\rho_A + \rho_B)^2 = 1 - \left[(\rho_A - \rho_B)/(\rho_A + \rho_B)\right]^2$ over the plane of final population sizes $\rho_A$ and $\rho_B$. Colour runs from blue at $\varphi = 0$ to dark red at $\varphi = 1$. Thin grey curves are iso-$\varphi$ contours, with the 0.6 and 0.8 levels labelled. The dashed line marks $\rho_A = \rho_B$, where $\varphi = 1$. All contours are straight rays through the origin, since $\varphi$ is scale-invariant and depends only on the ratio $r = \rho_A/\rho_B$ through $\varphi = 4r/(1 + r)^2$. The index therefore measures evenness alone and carries no information about total biomass. It falls to 0.8 at a two-fold imbalance and to 0.36 at a five-fold imbalance, and approaches 0 as either species goes extinct.

Table S2: Default model parameters. Values marked "varied" require the exact sweep values and replicate numbers in the archived simulation configuration or figure-specific reporting.

| Symbol | Description | Value |
|---|---|---|
| $\mu_{\max,A}$ | Mean maximum growth rate of species $A$ | 1.0 $\text{h}^{-1}$ |
| $\mu_{\max,B}$ | Mean maximum growth rate of species $B$ | 1.5 $\text{h}^{-1}$ |
| $\epsilon_\mu$ | Individual growth-rate variation | $\pm 20\%$ |
| $K_S$ | Substrate half-saturation constant | 0.5 a.u. |
| $K_M$ | Metabolite half-saturation constant | 0.5 a.u. |
| $\delta$ | Maintenance rate | 0.005 $\text{h}^{-1}$ |
| $Y_{X/S}$ | Biomass yield on substrate | 0.5 |
| $Y_{X/M}$ | Biomass yield on metabolite | 0.3 |
| $I$ | Metabolite-production investment | varied; 0.45 reference |
| $\lambda$ | Secretion efficiency | 10 |
| $n$ | Hill coefficient | 1 |
| $\theta$ | Hill activation threshold | $5 \times 10^{-4}$ a.u. |
| $S^\ddagger$ | Sequential-use switch threshold | 2.0 a.u. |
| $\beta_{\text{syn}}$ | Synergistic growth coefficient | 1.6 |
| $m^\ddagger$ | Division mass | 2.0 a.u. |
| $L_x \times L_y$ | Lattice dimensions | $51 \times 51$ patches |
| $\Delta x$ | Patch length | 10 $\mu$m |
| $D_S$ | Substrate diffusivity | 50 $\mu\text{m}^2\,\text{h}^{-1}$ |
| $D_M$ | Metabolite diffusivity | varied; 1 $\mu\text{m}^2\,\text{h}^{-1}$ reference |
| $k_d$ | Metabolite decay rate | varied; 0 $\text{h}^{-1}$ reference |
| $S_0$ | Initial substrate concentration | 100 a.u. |
| $\Delta t$ | Tick duration | 0.05 h |
| $N_0$ | Initial cells per species | 100 |
| $\ell$ | Separation in the reduced transport geometry | 5.8 patches |

### S6. Reduced metabolite-initiation threshold

The all-or-nothing establishment pattern of obligate bidirectional cross-feeding can be examined with a symmetric, well-mixed reduction. Assume $N_A = N_B = N_0$, equal cell mass $m$, a fixed and approximately saturating substrate response $f_S = S_0/(K_S + S_0)$, and identical metabolite pools. The gross rate of either obligate partner is

$$\mu_{\text{gross}}(M) = \mu_{\max} f_S \frac{M}{K_M + M} H(M). \tag{19}$$

Combining secretion (equation 10) with the ungated metabolite uptake in equation 12 gives

$$\frac{dM}{dt} = N_0 m \mu_{\max} \frac{M}{K_M + M} \left[ I \lambda f_S H(M) - \frac{1}{Y_{X/M}} \right]. \tag{20}$$

For $M > 0$, the sign of the metabolite feedback is determined by the bracketed term. Because $H(M) \to 0$ as $M \to 0$, the metabolite-free state is locally stable. A positive ignition threshold can exist only if

$$I \lambda f_S Y_{X/M} > 1. \tag{21}$$

The corresponding minimum investment is

$$I^* = \frac{1}{\lambda f_S Y_{X/M}}, \tag{22}$$

and, for $I > I^*$, the unstable metabolite threshold is

$$M^* = \frac{\theta}{\left( I \lambda f_S Y_{X/M} - 1 \right)^{1/n}}. \tag{23}$$

An initial metabolite concentration above $M^*$ produces positive metabolite feedback; a concentration below $M^*$ decays toward zero. With $\lambda = 10$, $Y_{X/M} = 0.3$, $S_0 = 100$ and $K_S = 0.5$, $f_S \simeq 0.995$ and equation 22 gives $I^* \simeq 0.335$.

This result is a metabolite-initiation condition, not by itself a proof of a population-level

Allee effect. Consortium establishment additionally requires

$$(1-I)\mu_{\max} f_S \frac{M}{K_M + M} H(M) > \delta \tag{24}$$

and depends on the initial metabolite seed. The derivation assumes symmetric partners, fixed substrate, fixed population size and well-mixed metabolite dynamics. It identifies the feedback responsible for the simulated threshold but is not expected to reproduce a spatial, stochastic transition exactly.

### S7. Obligate cross-feeding threshold ratio and optimal diffusivity

The transport result is approximated by a one-dimensional producer–receiver geometry. A producer at $x = 0$ releases metabolite at a fixed total rate $\sigma$, a receiver is located at $x = \ell$, and absorbing boundaries occur at $x = \pm L/2$. At steady state,

$$D\frac{d^2M}{dx^2} - k_d M + \sigma\delta(x) = 0, \qquad M(-L/2) = M(L/2) = 0. \tag{25}$$

Writing $\zeta = \sqrt{D/k_d}$, the solution is

$$M(x; D) = \frac{\sigma}{2\sqrt{Dk_d}} \frac{\sinh[(L/2 - |x|)/\zeta]}{\cosh[L/(2\zeta)]}. \tag{26}$$

To retain the implemented obligate growth kinetics, define $M_{\text{crit}}$ implicitly by

$$(1-I)\mu_{\max,r} f_{S,r} \frac{M_{\text{crit}}}{K_{M,r} + M_{\text{crit}}} H(M_{\text{crit}}) = \delta_r. \tag{27}$$

The dimensionless cross-feeding threshold ratio used in the main text is then

$$R_0(D) = \frac{M(\ell; D)}{M_{\text{crit}}}. \tag{28}$$

Under the assumptions of this reduction, $R_0 > 1$ indicates that delivered metabolite exceeds the concentration required for positive receiver growth, whereas $R_0 < 1$ indicates that it does not. This quantity is a metabolite sufficiency ratio; it is not a next-generation reproduction number in the epidemiological sense.

For a domain large relative to the reaction–diffusion length ($L \gg \zeta$), equation 28 becomes

$$R_0(D) \simeq C D^{-1/2} \exp\left(-\ell\sqrt{k_d/D}\right), \qquad C = \frac{\sigma}{2M_{\text{crit}}\sqrt{k_d}}. \tag{29}$$

Maximizing this approximation gives

$$D^* = \ell^2 k_d, \qquad \sqrt{D^*/k_d} = \ell. \tag{30}$$

The approximation predicts that metabolite transport is maximized at the receiver when the reaction–diffusion length matches partner separation.

Equation 30 requires $k_d > 0$, $L \gg \zeta$, a fixed source strength $\sigma$ and receiver kinetics independent of $D$. For a finite absorbing domain, the optimum should be obtained from the exact expression in equation 26. If $k_d = 0$, the large-domain approximation and equation 30 are not applicable; the finite-domain boundary-loss problem must be solved directly. These conditions limit the analytical result to a mechanistic approximation of the simulation rather than a parameter-free prediction.
